# Evaluation of Anti-Fibrotic Therapeutics Using a Three-Dimensional *In Vitro* Liver Fibrosis Model

**DOI:** 10.1101/2025.11.24.690128

**Authors:** Mrunmayi Gadre, Kirthanashri S Vasanthan

## Abstract

Three dimensional (3D) bioprinting is a leading technology in tissue engineering that offers controlled deposition of cells and bioinks layer by layer, this technique allows for the accurate replication of tissue spatial organization, incorporating tissue compartmentalization and vascularization. This study focuses on the screening of anti-fibrotic drug on a previously established 3D liver diseased model. In market, there are no available 3D *in vitro* liver disease model that is used for liver regeneration/drug screening. The 3D *in vitro* liver disease was fabricated by utilizing decellularized rat liver Extracellular Matrix (dECM) and Gelatin Methacryl (GelMA) along with hepatic cells. The developed healthy model was rendered fibrotic by employing methotrexate (MTX) a fibrotic agent at a concentration of 10mM for 72 hours. MTX is well known to cause hepatotoxicity and have been analysed for causing fibrotic -like characteristic in the model. This manuscript mainly focuses on the reversal aspect of the study, where anti-fibrotic drug, aspirin was administered on the fibrotic model to evaluate the effect. This study examines the impact of antifibrotic drug aspirin (ASP) on both 2D HepG2 cell and the fabricated 3D model by integrating major experiments including the biochemical, morphological, and molecular analyses. The outcomes disclose a progressive decline in the fibrotic trait and regaining the hepatocyte-specific functionality. Successful employment of this 3D model will reduce the involvement of animals in drug testing experiments.

## 1. INTRODUCTION

Liver fibrosis represents a critical health challenge globally, characterized by the accumulation of extracellular matrix proteins in excessive amounts, that includes collagen, in response to chronic liver injury [1]. This pathological condition serves as a precursor to cirrhosis, a major contributor to liver-related mortality. Liver fibrosis is the wound-healing response by the liver to persistent damage, such as chronic viral infections, alcohol abuse, non-alcoholic fatty liver disease and autoimmune disorders [2]. Liver fibrosis is reversible; however, as persistence of injurious stimuli, it progresses to cirrhosis, leading to hepatocellular carcinoma and eventually failure [3,4]. The progression of fibrosis is staged from F0 (no fibrosis) to F4 (cirrhosis), with intermediate stages indicating varying degrees of scarring [5]. **Figure 1** simplifies the understanding and the differences in all stages between F0 to F4. In spite of the fact that liver fibrosis remains a global health burden, there is no effective anti-fibrotic drugs available.

**Figure 1:**
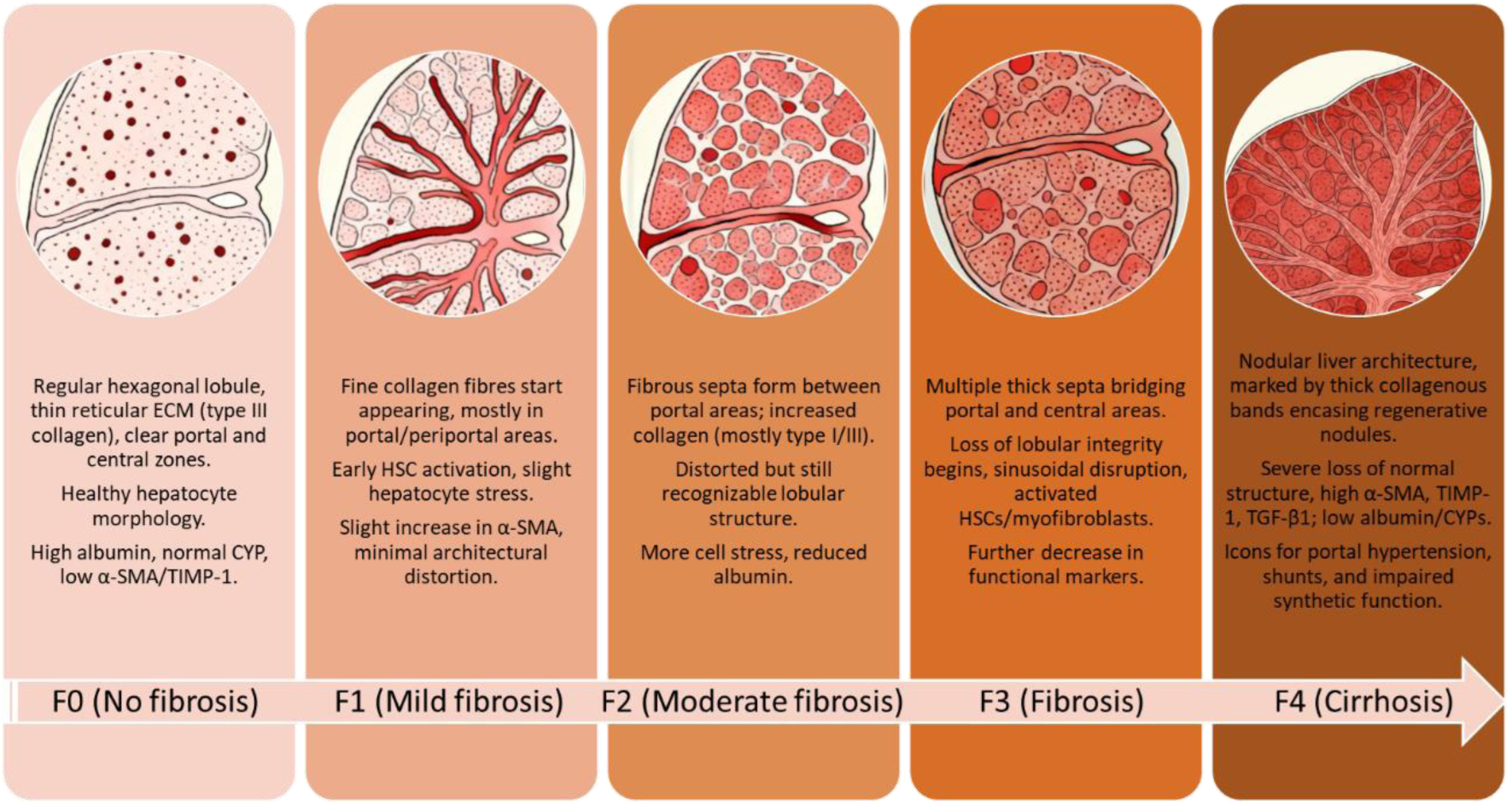
Differences stages of fibrosis (F0 to F4)

At present the only therapy offered for curing liver fibrosis is aiming to control underlying causes of liver injury. The complex pathways and signaling cascades involved in fibrosis and reversal make it difficult for any anti-fibrotic drug to effectively reverse fibrosis and pass clinical trials. Another major challenge also occurs when there is a lack of presence of a physiologically relevant *in vitro* model [6]. Although the traditional two-dimensional (2D) cell culture is utilized for the initial drug screening, they fail to replicate the multicellular interactions, extracellular matrix (ECM) composition, and mechanical cues of the native liver microenvironment [7]. On the contrary, the animal models utilised for the screening of drugs may lead to poor translation of preclinical results into clinical success as they do not fully mimic human fibrotic responses. This major gap can be overcome with the application of 3D *in vitro* liver models; these platforms allow the replication of disease conditions and therapeutic screening under human-relevant conditions [8]. These models enable in the maintaining of cellular heterogeneity, mimic the liver’s microarchitecture, and support the study of disease progression and drug responses.

Current methods in the pharmaceutical industry for testing anti-liver fibrosis drugs primarily depend on two-dimensional (2D) cell culture systems because they are straightforward, cost-efficient, and compatible with high-throughput screening [9]. The standard models often utilize hepatic specific cell lines such as primary hepatic stellate cells or LX-2 with evaluations centred on fibrosis-associated markers like α-SMA and collagen production [10, 11]. The rigidity and flatness of traditional culture plastics do not match the biomechanical properties of fibrotic liver tissue, often causing inaccuracies in predicting a drug’s real-world efficacy and safety. As a result, compounds that initially seem effective in 2D assays frequently fail or show reduced performance when moved into more complex three-dimensional or animal models, highlighting a translational gap and the urgent need for new systems that better reflect the *in vivo* fibrotic liver environment [12].

The goal of this study is to overcome the current limitation of drug screening by integrating Gelatin methacylamide (GelMA), a material known for its robust mechanical stability along with rat liver-derived decellularized extracellular matrix (dECM), which provides essential native biochemical signals [13]. These two components along with HepG2 cells and crosslinking agents forms a bioink that creates environment that supports hepatic cell function. Post 3D bioprinting, the construct closely mimics the natural structure and activity of healthy and fibrotic liver tissue and has been established in previous study by the group [14].

Figure 2 is the overall methodology used for this study. This 3D model is not only better able to replicate the native physiology and pathology of liver tissue, but also makes the modelling of diseases, screening of drugs and toxicity testing more accurate and reliable. Ultimately, this approach shows great potential for accelerating the discovery and validation of new therapies for liver diseases, as it provides a highly relevant and translational platform for preclinical studies. The integration of 3D in vitro liver models into research and drug development pipelines holds great promise for advancing our understanding of liver fibrosis and improving therapeutic outcome. By bridging the gap between traditional 2D cultures and in vivo studies, these models provide a more accurate and ethical way of studying liver diseases and testing potential treatments. Continued innovation and refinement of 3D liver models are necessary for the translation of laboratory findings into clinical applications, which will ultimately result in improved management and treatment of liver fibrosis.

**Figure 2:**
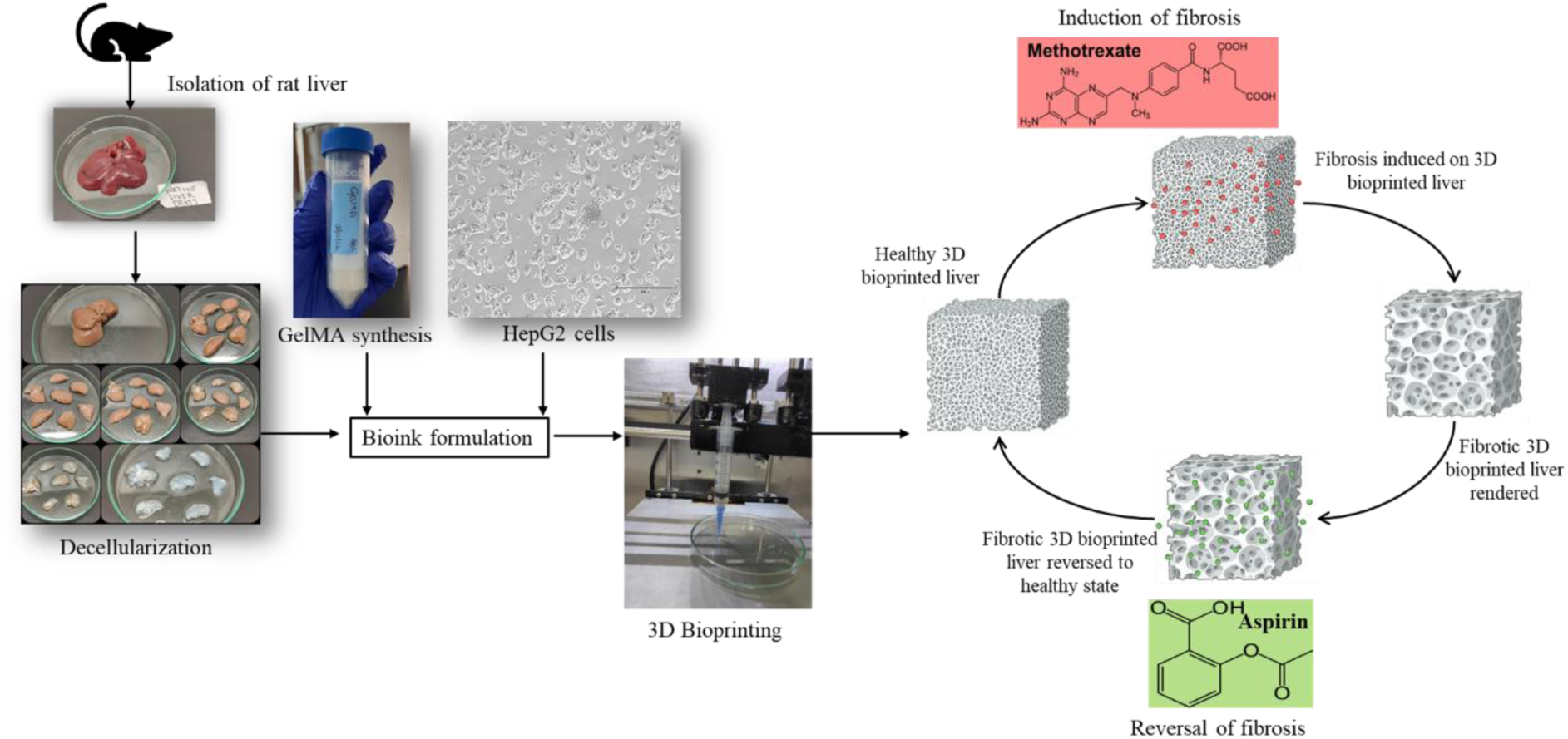
Overview flowchart of the study

### 2.0 MATERIALS AND METHODOLOGY

### 2.1 Materials

The cell culture experiments were conducted using Dulbecco’s Modified Eagle’s Medium (DMEM) with high glucose (4.5g/L), L-glutamine, and sodium bicarbonate, but without sodium pyruvate. DMEM and Dulbecco’s Phosphate-Buffered Saline (DPBS) were sourced from Himedia, India. Fetal bovine serum (FBS) was supplied by Gibco. Essential reagents, including penicillin/streptomycin, trypsin, trizol, and methotrexate, were purchased from Sigma Aldrich. The Albumin (BCG) Assay Kit (ab235628) came from Abcam, while lactate dehydrogenase (LDH) activity assay [E-BC-K046-M] and urea (BUN) colorimetric assay [E-BC-K183-M] were acquired from Elab Sciences.

For enzymatic activity assessments, alanine transaminase (ALT) [ab241035] and alkaline phosphatase (ALP) [ab83369] colorimetric assay kits were obtained from Abcam. Chemicals such as sodium dodecyl sulfate, phenol, chloroform, isoamyl alcohol, isopropanol, and papain were purchased from Sisco Research Laboratories, India. Ethylenediamine tetraacetic acid (EDTA) and formaldehyde came from Qualigens, India, and glutaraldehyde from Sigma Aldrich, India. Proteinase K was supplied by Invitrogen, India, while Coomassie Brilliant Blue dye and a protease inhibitor cocktail were procured from Merck, India. Sodium acetate was sourced from Tokyo Chemical Industry, India.

Additional reagents included RIPA buffer from Himedia, India; Pierce™ CA Protein Assay Kit from Thermo Fisher Scientific, India; Blyscan™ sulfated glycosaminoglycan (sGAG) assay kit from Biocolor, United Kingdom; and both the PrimeScript™ 1st Strand cDNA Synthesis Kit and TB Green Premix Ex Taq (Tli RNase H Plus) from Takara, USA.

### 2.2 Animal ethics approval

All the protocols related to the animals in this study was performed with approval of Institutional Animal Ethics Committee (IAEC), protocol number: IAEC/KMC/121/2022, at Manipal Academy of Higher Education, Manipal. The guidelines are aligned with the Committee for the Purpose of Control and Supervision of Experiments on Animals (CPCSEA). Under strict adherence to the ARRIVE guidelines, all the experiments involving animals were carried out to ensure ethical and humane treatment throughout the research process.

### 2.3 METHODOLOGY

#### 2.3.1. The Cell Culture technique

The Human hepatocellular carcinoma cell culture (HepG2) was procured from National Centre for Cell Science (NCCS), Pune, India. Cells under complete culture conditions were cultured in Dulbecco’s Modified Eagle Medium (DMEM) which was supplemented with 10% (v/v) fetal bovine serum (FBS) along with 1% (v/v) penicillin/streptomycin. The cells were sustained in a humidified incubator at 37°C with 5% COL to ensure optimal growth conditions.

#### 2.3.2. Rat Liver Harvest and Chemical Decellularization

Male Wistar rats (9-12 weeks of age and 150-250 grams) were first anesthetized with an intraperitoneal injection of sodium pentobarbital at a concentration of 50 mg/kg. Following anesthesia, livers were carefully excised and thoroughly rinsed in sterile distilled water and placed in 0.001 M ethylenediaminetetraacetic acid (EDTA) solution for 30 minutes to promote initial matrix slackening. Post this, the liver was cut into small pieces and then perfused with approximately 60 mL of double distilled water to ensure that all traces of blood were removed. Decellularization of liver was performed through the successive perfusion of sodium dodecyl sulfate (SDS) with increase in concentrations, i.e., 0.1%, 0.25%, 0.5%, 0.75%, and 1%, (each 60-80 mL), through 7-11 sites of injection per tissue segments, ensuring even distribution and coverage. After performing the perfusion procedures, the tissues were thoroughly rinsed with distilled water until they were translucent. This indicates decellularization was successful based on visual confirmation. The obtained decellularized extracellular matrix (dECM) was further characterized regarding morphological and biochemical properties.

#### 2.3.3. Synthesis of Gelatin Methacryloyl (GelMA)

GelMA was synthesis using a modified protocol reported in the literature [13]. Briefly, the type A gelatin (Bloom strength 175 g, porcine source) was dissolved in PBS (pH 7.4) at 10% (w/v) and kept at 50°C until completely dissolved. Then, the crosslinker Methacrylic anhydride (MA) was slowly added (0.6 g MA per 1 g gelatin) with continuous stirring (1200 rpm) at 50°C and then allowed for reaction for 60 mins. The mixture was first centrifuged at 4000 rpm for 2 minutes that helps remove the excess MA. The pellet was discarded and equal amount of 1X PBS was added to the supernatant. The diluted mixture was dialyzed against deionized water using a 10 kDa molecular weight cut off (MWCO) dialysis membrane with and kept at a temperature of 40°C for five days with daily water changes to remove the unreacted content. The obtained GelMA was lyophilized and stored at -20°C until further use. Characterization of GelMA involves comparison with native gelatin physicochemical properties.

#### 2.3.4. Computer-Aided Design (CAD) for 3D Bioprinting

CAD required for 3D bioprinting were created applying Openscad and exported as stereolithography (STL) files. The bioprinting specifications specific to the model that includes geometric dimensions and extrusion parameters were precisely defined in the software environment. CAD designs were generated either from medical imaging modalities (X-ray, CT, MRI) or constructed directly using the design software. All STL files were then validated using the computer-aided manufacturing (CAM) tools that confirmed the printability and compatibility with the chosen bioprinting system, and extrusion volumes of bioink weas calculated based on intended layer dimensions in the STL files.

#### 2.3.5. Bioink Preparation and Bioprinting

A photocrosslinkable and enzyme curable GelMA-based bioink was made by reconstituting the GelMA that was lyophilized in PBS with lithium phenyl-2,4,6-trimethylbenzoylphosphinate (LAP; Sigma-Aldrich, China) and microbial transglutaminase (MTgase) and homogenized with HepG2 cells. The resulting cell-laden bioink was filled in sterile syringe (0.7 mm inner diameter needle) and dispensed with a dual-extrusion bioprinter (Alfatek Systems, India). Bioprinting was perfomed at room temperature with pressure of 120-180 kPa and a speed of 600 mm min-1 to obtain 1 x 1 cm constructs in a rectilinear lattice architecture. Crosslinking was performed by short UV irradiation in the presence of LAP in order to stabilize the printed GelMA constructs.

#### 2.3.6. Induction of Fibrosis

Fibrosis was induced in HepG2 cell cultures (both on standard 2D plates and 3D bioprinted constructs) by treating them with 10 mM methotrexate over a 72 hour period. Methotrexate concentrations and exposure times were selected based on prior optimization studies. Post treatment period, the cell viability performed using the MTT assay, and morphological evaluation of fibrotic changes was performed by haematoxylin and eosin (H&E) staining. For both the 3D cultured and 3D bioprinted scaffolds, the cells were fixed in formaldehyde, dehydrated successively increasing graded ethanol wash, then stained, and finally mounted with DPX for analysis.

#### 2.3.7. Reversal of Fibrosis

For the experiments studying the reversibility of fibrosis, aspirin was used in both 2D HepG2 cultures and HepG2 cultivated on 3D bioprinted constructs. The concentration and treatment duration of aspirin was optimized in 2D cultures first before use on 3D bioprinted tissues. Following aspirin exposure, the cell viability measured by MTT assay along with morphological changes was observed by H&E staining using the same fixation, dehydration, staining and mounting protocol described previously.

#### 2.3.8. Characterization of 2D Cell Culture and 3D Constructs for Anti-Fibrotic Drug Screening

##### 2.3.8.1. Cell Viability Assay

Cell viability was evaluated by adding 0.5% MTT solution to the wells (100 μL per well in 96-well plates) and incubating for 2 hours at 37°C. After removal of MTT, to dissolve the formazan crystals 100 μL of DMSO was added. At three different absorbance wavelength was measured including 570 nm, 590 nm, and 630 nm with a microplate reader to determine cell metabolic activity.

##### 2.3.8.2. Live/Dead Assay

The assay was performed on HepG2 cells following fibrosis induction, strictly rendering to instructions provided by the manufacturer. Briefly, a solution was prepared, diluting 0.2 μM calcein AM in 1 mL DPBS and 0.4 μM EthD-1 in the same solution. Culture media was discarded, and cells were briefly washed with DPBS before exposure to 100 μL of the working solution per well (96-well plate). Under light-blocked conditions incubation was performed for 45 minutes, after which the stained cells were observed using an EVOS M5000 fluorescence microscope (Invitrogen, India). From this experiments onward as the concentration of 2mM ASP was proven to be optimum and used to reverse the fibrosis on the 3D scaffolds on day 5.

##### 2.3.8.3. Histology and Ishak scoring

For histological assessment, cells were first washed twice with 1X PBS and fixed using 4% ice-cold formaldehyde for 15 minutes. Post-fixation, cells were washed three times in PBS and permeabilized with 0.2% Triton X-100 for 20 minutes. Hematoxylin staining was performed for 1 minute at room temperature, followed by running tap water rinse and 45 seconds of eosin staining. Rapid washes were done sequentially in 95% and 100% ethanol before final mounting in DPX for microscopic visualization under the EVOS M5000. The slides were then outsourced for ishak scoring and were scored according to the standard [15].

##### 2.3.8.4. Efflux Media Collection and Biochemical Measurements

Spent culture media were collected from healthy 3D bioprinted models on days 1, 3, 5, and 7, and from fibrotic models on days 1, 2, and 3. Key liver enzyme concentrations—alkaline phosphatase (ALP) and alanine transaminase (ALT)—were measured using commercially available kit-based methods as per manufacturer’s instructions.

###### 2.3.8.4.1. Albumin Content

Albumin concentration was determined via a colorimetric kit assay utilizing Bromocresol Green (BCG), which forms a measurable chromophore with albumin at 620 nm. For the analysis, 50 μL of sample was added to kit reagent (100 μL), 25 minutes at room temperature was incubation time, and absorbance recorded at 620 nm. Albumin concentrations were then calculated with the reference of standard curve.

###### 2.3.8.4.2. Urea Content

The levels of urea were quantified by a calorimetric kit based method. In brief, 4 μL of sample was mixed with enzyme solution (50 μL) and incubated for 10 minutes at 37°C. Subsequently, reagents 4 and 5 (125 μL each) were added, followed by further incubation 10 minute. Urea concentrations were then calculated with the reference of standard curve.

###### 2.3.8.4.3. ALP Content

ALP activity were again assessed by colorimetric kit (Abcam), using p-nitrophenyl phosphate (pNPP) as substrate. Wells received 50 μL of 5 mM pNPP solution, followed by 10 μL of ALP enzyme for standard wells, and incubation at 25°C for 60 minutes. Reactions were terminated by 20 μL stop solution per well, and ALP concentrations were then calculated with the reference of standard curve.

###### 2.3.8.4.4. ALT Content

ALT was measured by a rapid colorimetric kit (Abcam). Samples (0.5-2.5 mL, made up to 5mL with buffer if needed) were added to wells (25mL freshly prepared reaction mix and 25mL background mix for controls). Absorbance was read at 570 nm after cycles of kit and the ALT levels were extrapolated using a control standard curve.

###### 2.3.8.4.5. LDH Content

Lactate dehydrogenase (LDH) was assayed by colorimetric method constructed on the reaction of pyruvate dinitrophenylhydrazone and lactic acid to pyruvic acid and to yield brown-red color in alkaline solution. 200 mL sample, 50 mL coenzyme I and 250 mL substrate buffer were incubated at 37°C for 15 min and 250 mL chromogenic agent and 10 min of incubation were applied. Alkali application solution (2500 mL) was then added, and LDH concentrations were then calculated with the reference of standard curve at 450 nm.

##### 2.3.8.5. Quantitative Real-Time RT-PCR

Complete RNA was first extracted from treated cell pellets using the standard TRIzol-based protocol. Briefly, pellets were washed once with 1x PBS and lysed in 800 uL TRIzol reagent. Phase separation was induced by adding 160 uL chloroform, followed by vigorous vortexing and incubation for 2 min at room temperature. Contents were then centrifuged for 20 min at 4°C at 12,000 rpm. The RNA which is in the aqueous phase was transferred to fresh tubes, mixed with an equal volume of molecular-grade isopropanol, vortexed, and allowed to stand at room temperature for 10 min to facilitate RNA precipitation. The resulting RNA was pelleted by centrifugation for 20 min at 4 °C at 13,000 rpm, washed twice with 1 mL of ice-cold 70% ethanol (centrifugation at 7,500 rpm for 5 min at 4 °C each time), and briefly air-dried on sterile tissue. Pellets were finally dissolved in 10 uL DEPC-treated water. RNA concentration and purity of RNA were determined spectrophotometrically further processed for cDNA synthesis.

For first-strand cDNA synthesis, kit components were thawed on ice and a reaction master mix was prepared according to the manufacturer’s instructions. A volume of 7 uL of the master mix was dispensed into each tube, to which the required amount of RNA and nuclease-free water were added to achieve the recommended reaction volume and RNA input. Reverse transcription was carried out with the following program: 37 °C for 30 min, 85°C for 5 s to inactivate the reverse transcriptase, followed by a hold at 4 °C. cDNA products were stored at -20 °C until quantitative RT-PCR analysis. For RT-PCR, reaction mixtures and primer cocktails were assembled as per kit specifications; 6 uL of primer mix and 4 uL of either cDNA template or nuclease-free water (non-template control) were added to each reaction. Amplification was performed under the following cycling conditions: initial denaturation at 95 °C for 30 s (1 cycle); 40 cycles of 95 °C for 5 s and 60 °C for 34 s; followed by 95 °C for 15 s, 60 °C for 1 min, and 95 °C for 15 s for melt-curve analysis. Amplicons were stored at [?]20 °C until further analysis. The following primers were used:

##### 2.3.8.5. Quantitative Real-Time RT-PCR

- GAPDH forward: ACAGTCAGCCGCATCTTCTT
- GAPDH reverse: GACAAGCTTCCCGTTCTCAG
- Albumin forward: TGCTAATTTCCCTCCGTTTG
- Albumin reverse: CTGAGCAAAGGCAATCAACA
- Collagen I forward: ATGCCTGGTGAACGTGGTAGGAGAGCCATCAGCACCT
- Collagen I reverse: ACCAGCATCACCTCTGTCACCCTT

## 3.0 RESULTS AND DISCUSSION

### 3.1. Fabrication of 3D Bioprinted Construct

#### 3.1.1 Rat liver decellularization

Rat livers were decellularized using a sequential chemical–enzymatic regimen that depleted cellular material while preserving extracellular matrix (ECM) architecture. H&E staining showed loss of nuclei, consistent with efficient cell clearance. Tissue mass declined by ∼87%, indicating removal of cellular components. SEM distinguished compact hepatocytic organization in native tissue from the porous, fibrous network characteristic of decellularized scaffolds. DNA content dropped from ∼1200 ng/mg (native) to ∼10 ng/mg (dECM); agarose gels showed no residual genomic bands. SDS-PAGE demonstrated retention of major ECM proteins (e.g., collagens/glycoproteins), supporting matrix integrity and low immunogenicity. These outcomes satisfy accepted benchmarks for effective decellularization (≤50 ng DNA/mg dry tissue with preserved ECM proteins). The dECM was then solubilized for homogeneous incorporation into the bioink.

#### 3.1.2 GelMA synthesis

Gelatin methacryloyl (GelMA) was prepared via a modified methacrylation to achieve efficient substitution of amino groups for photocrosslinking. FTIR revealed the expected methacrylate signatures, and ^1H-NMR confirmed successful functionalization. Expanded physicochemical and rheological properties of this GelMA batch are reported in our prior patent filing [13], supporting its mechanical robustness and biocompatibility.

#### 3.1.3 Three-dimensional bioprinting

CAD models were converted to G-code for extrusion printing of rectilinear lattices (≈1 × 1 × 0.3 cm³) with interconnected porosity to aid perfusion and cell survival. A hybrid bioink—GelMA + solubilized rat liver dECM + HepG2 cells—yielded reproducible, mechanically stable constructs. The tunable mechanics of GelMA combined with organ-specific biochemical cues from dECM produced a microenvironment compatible with hepatocyte function. Prior work indicates that liver-derived ECM in GelMA enhances hepatocyte adhesion, polarity, and albumin production, positioning these constructs as a suitable platform for liver fibrosis modeling and anti-fibrotic drug evaluation.

### 3.2 Induction of fibrosis

Fibrosis-like phenotypes were induced by 10 mM methotrexate (MTX) for 72 h, as optimized previously [14]. MTX is a well-known hepatotoxicant that promotes oxidative stress and matrix accumulation, recapitulating features of fibrogenesis. Post-treatment microscopy showed increased cell density, disruption of hepatic architecture, and collagenous deposition, confirming successful induction. Unlike 2D monolayers, the 3D GelMA–dECM system preserves cell–ECM and mechano-biological cues, enabling more physiomimetic progression of fibrotic changes. This controllable model therefore bridges the gap between simplified in vitro assays and in vivo studies and supports more reliable preclinical screening of anti-fibrotic candidates.

### 3.3. Characterization of 2D Cell Culture and 3D Constructs for Anti-Fibrotic Drug Screening

#### 3.3.1 Cell Viability Assay

Cell viability was evaluated in 2D monolayers and 3D bioprinted fibrotic constructs at Day 3 (post-fibrosis induction) and again at Day 5 after anti-fibrotic treatment. Pre-assay phase-contrast images **(Figure 3)** showed a progressive reduction in cell density and loss of typical hepatocyte morphology with increasing aspirin (ASP) concentration, consistent with drug-driven cytotoxicity at higher doses.

**Figure 3:**
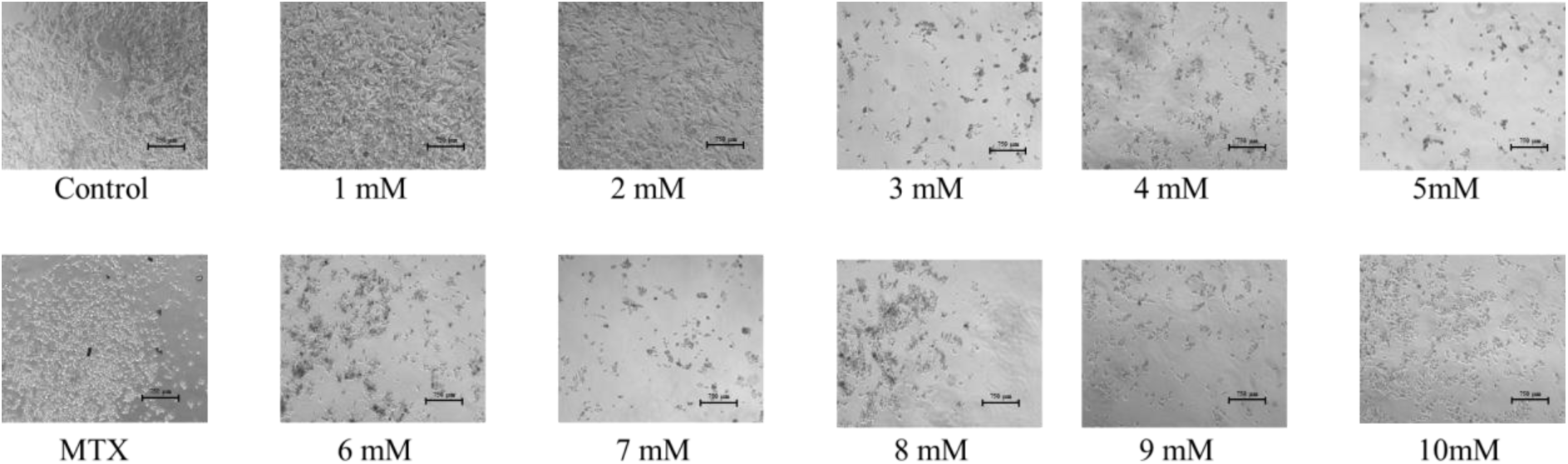
Microscopic images of HepG2 cells post treatment with aspirin

MTT quantification **(Figure 4A)** mirrored these observations: metabolic activity declined in a clear dose-dependent manner across the ASP gradient, indicating an inverse relationship between viability and ASP concentration. Based on this profile, 2 mM ASP was selected as the working dose for 3D studies, balancing anti-fibrotic efficacy with acceptable cytocompatibility.

**Figure 4:**
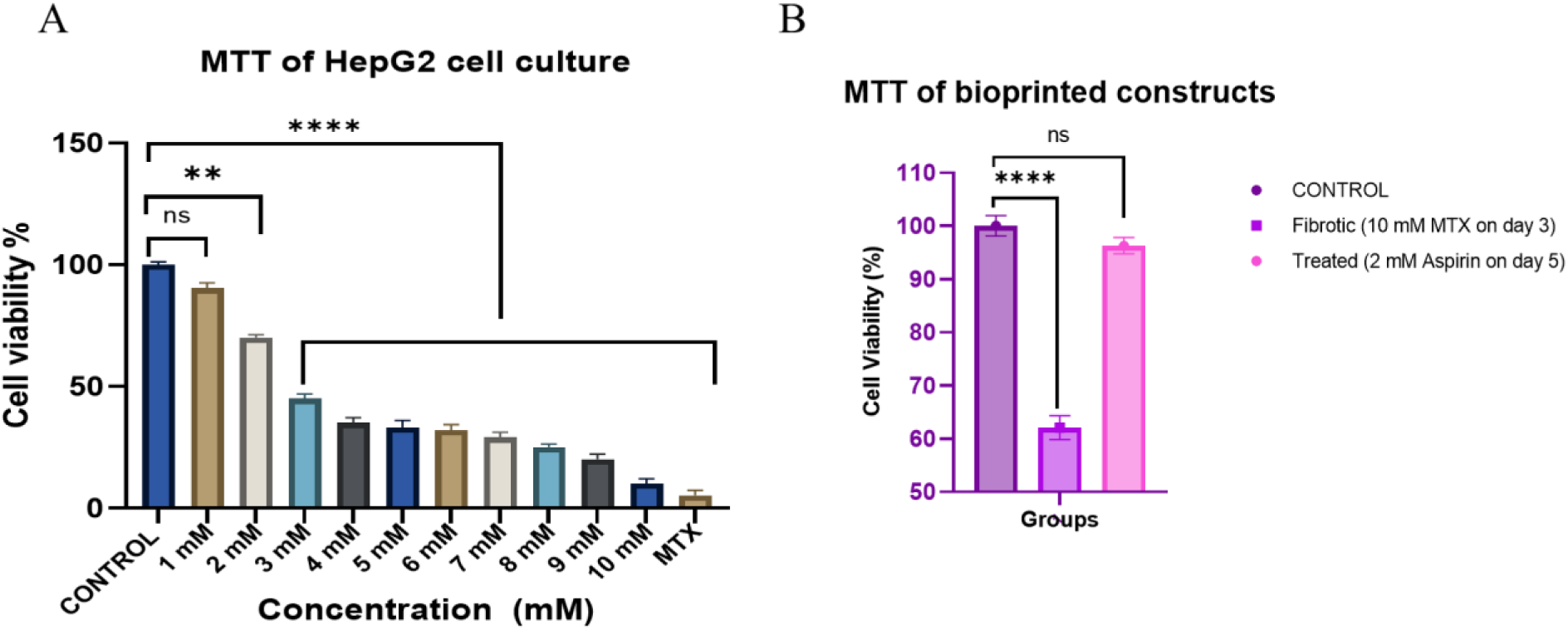
Cell viability analysis via MTT A: 2D culture and B. 3D scaffolds; statistically significant ** : p<0.01, **** : p< 0.0001 and ns: non-significant

Applying 2 mM ASP to the fibrotic models **(Figure 4B)** showed that post-treatment viability in both 2D and 3D groups remained statistically comparable to their respective controls (p > 0.05), indicating that the chosen dose does not compromise cell survival while enabling fibrosis reversal experiments. Notably, the 3D GelMA–dECM constructs-maintained viability at least on par with 2D cultures after treatment, consistent with the protective, physio mimetic microenvironment of the printed matrix. Collectively, these data validate 2 mM ASP as a suitable concentration for downstream anti-fibrotic assessments in the 3D liver model.

#### 3.3.2 Live/Dead Assay

Live/Dead staining was performed on 2D monolayers and 3D bioprinted fibrotic constructs at Day 3 (after MTX-induced fibrosis) and at Day 5 following anti-fibrotic treatment. Representative micrographs (**Figure 5; calcein AM, green = live; EthD-1, red = dead; scale bar: 750 µm)** show a dose-dependent decline in the live cell fraction with increasing ASP, accompanied by a corresponding rise in red fluorescence.

**Figure 5:**
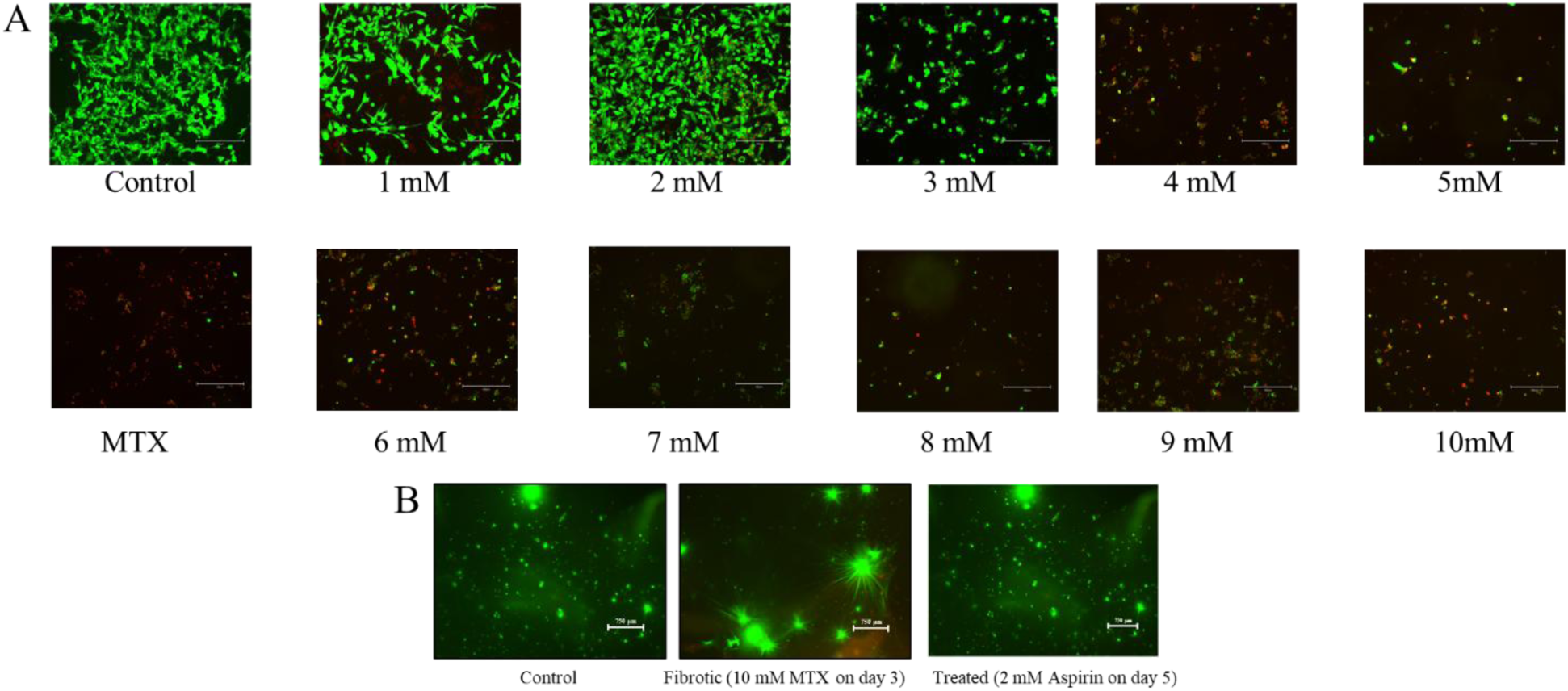
Cell viability analysis via Live/dead assay A: 2D culture and B. 3D scaffolds

Consistent with the MTT results, viability was well maintained at 2 mM ASP, whereas concentrations above 2 mM produced a marked loss of green signal and increased EthD-1 uptake, indicating cytotoxicity. After reversal on Day 5 with 2 mM ASP, both 2D and 3D groups displayed robust green fluorescence relative to untreated fibrotic controls, with the 3D GelMA–dECM constructs retaining a higher proportion of live cells than 2D cultures at equivalent treatment. These observations support 2 mM ASP as an efficacious yet cytocompatible working concentration for subsequent anti-fibrotic evaluations in the 3D liver model.

#### 3.3.3 Histology and Ishak Scoring

H&E staining of the bioprinted constructs **(Figure 6; scale bar 75 um)** revealed obvious morphological changes correlated with the induction of fibrosis and reversal of fibrosis by pharmacology. After MTX exposure, the parenchyma looked disorganized with higher cell density and spindle-shaped, myofibroblast-like cells (circled), consistent with a contractile phenotype and tissue remodeling. These features were associated with a higher Ishak score (grade 4) determined by blinded evaluation, suggesting significant fibrotic change in the 3D matrix. Following treatment with 2 mM ASP, the constructs showed recovery towards normal hepatic morphology: cells showed their round/epithelioid appearance back, intercellular spaces were more uniform, and the diffuse condensation of stroma observed in MTX-treated samples was reduced. Correspondingly the Ishak score improved to grade 1, indicating marked attenuation of fibrosis. Together with the MTT and Live/Dead data, these histological results support the idea that ASP at 2 mM is able to elicit anti-fibrotic effects without affecting viability. The 3D GelMA-dECM microenvironment allows one to visualize structural regression (cell shape, tissue organization) that is hard to capture in 2D, highlighting the translational relevance of this platform for phenotypic readouts and quantitative scoring of fibrosis reversal.

**Figure 6:**
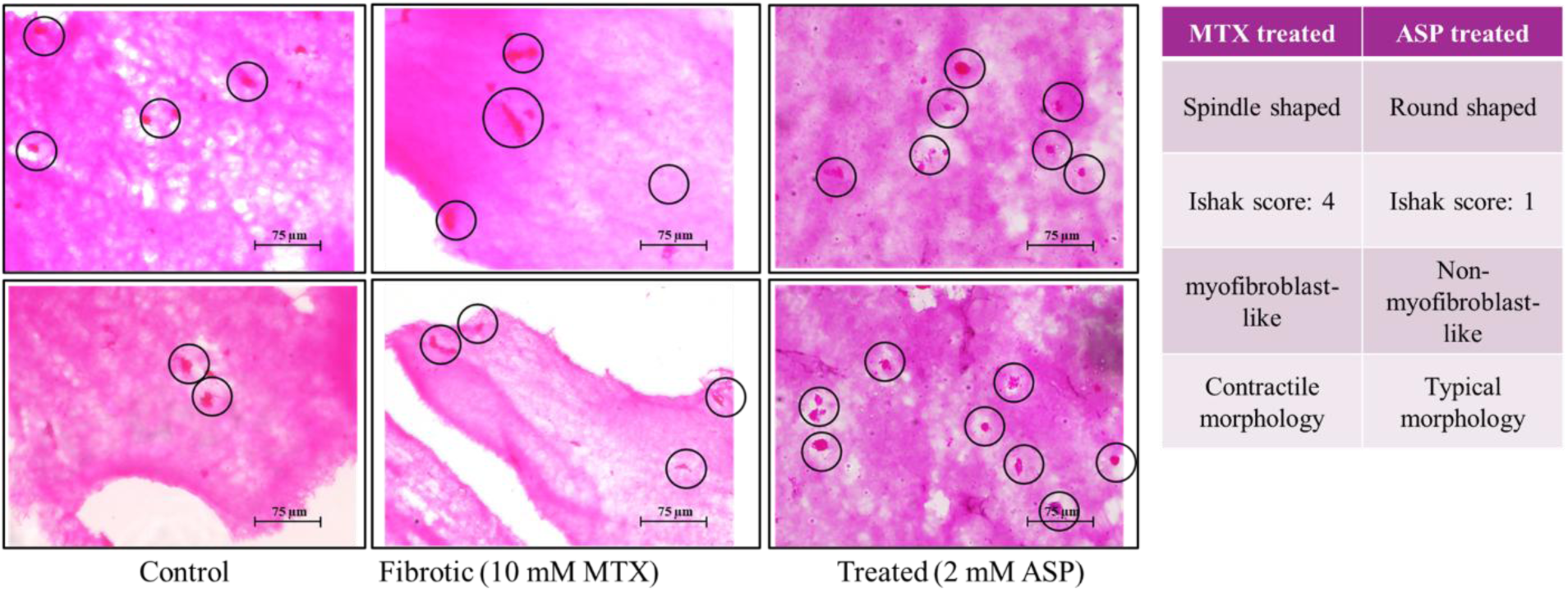
Histological analysis via H&E staining

#### 3.3.4 Efflux Media Collection and Biochemical Measurements

Liver-specific function of the bioprinted constructs was evaluated by assessing secretory (albumin, urea) and injury/leakage markers (LDH, ALP, ALT) in the spent media in control (non-fibrotic), MTX-fibrotic and ASP-treated conditions (**Figure 7**). As expected for a healthy hepatocytic phenotype, control constructs secreted more albumin and urea than MTX treated constructs, suggesting the protein synthesis and urea-cycle activity were preserved. In contrast, fibrotic constructs showed much higher LDH, ALP and ALT levels, which are consistent with membrane damage (LDH release) and hepatocellular/cholestatic injury (ALT/ALP). After anti-fibrotic treatment with 2 mM ASP, albumin and urea levels returned toward control values and LDH, ALP, and ALT levels decreased, so differences between ASP-treated and control groups were not statistically significant (p > 0.05) at the assay time point. Taken together these trends show that ASP not only alleviates the fibrotic morphology (Sections 3.3.2-3.3.3) but also restores hepatic functionality in the 3D GelMA-dECM constructs, strengthening the translational potential of this platform for functional readouts of fibrosis reversal.

**Figure 6:**
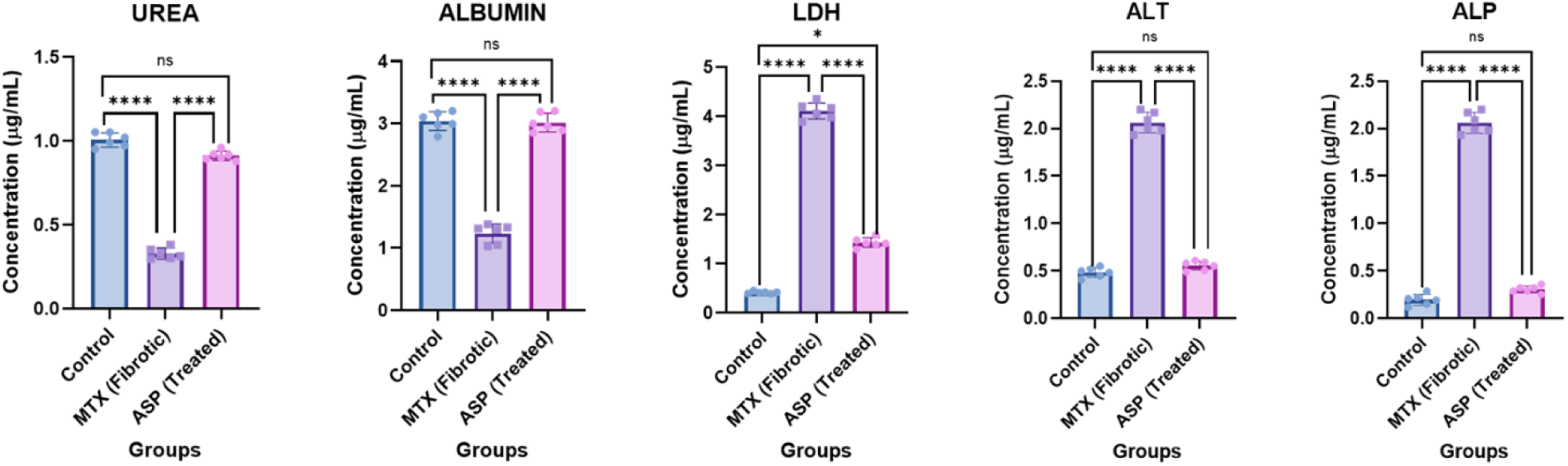
Functional assays performed on the 3D scaffolds; statistically significant **** : p< 0.0001 and ns: non-significant

#### 3.3.5 Quantitative Real-Time RT-PCR

Gene expression was profiled in control (non-fibrotic), MTX-fibrotic, and ASP-treated 3D constructs. Ct values were normalized to GAPDH and expressed as relative fold change (ΔΔCt) **(Figure 8)**. Hepatocyte function. The hepatocyte marker albumin (ALB) was significantly downregulated in MTX-treated constructs relative to control, consistent with loss of hepatic function under fibrotic stress. Following treatment with 2 mM ASP, ALB expression rebounded toward control, with differences no longer significant versus control at the assayed time point, indicating restoration of functional phenotype.

**Figure 8:**
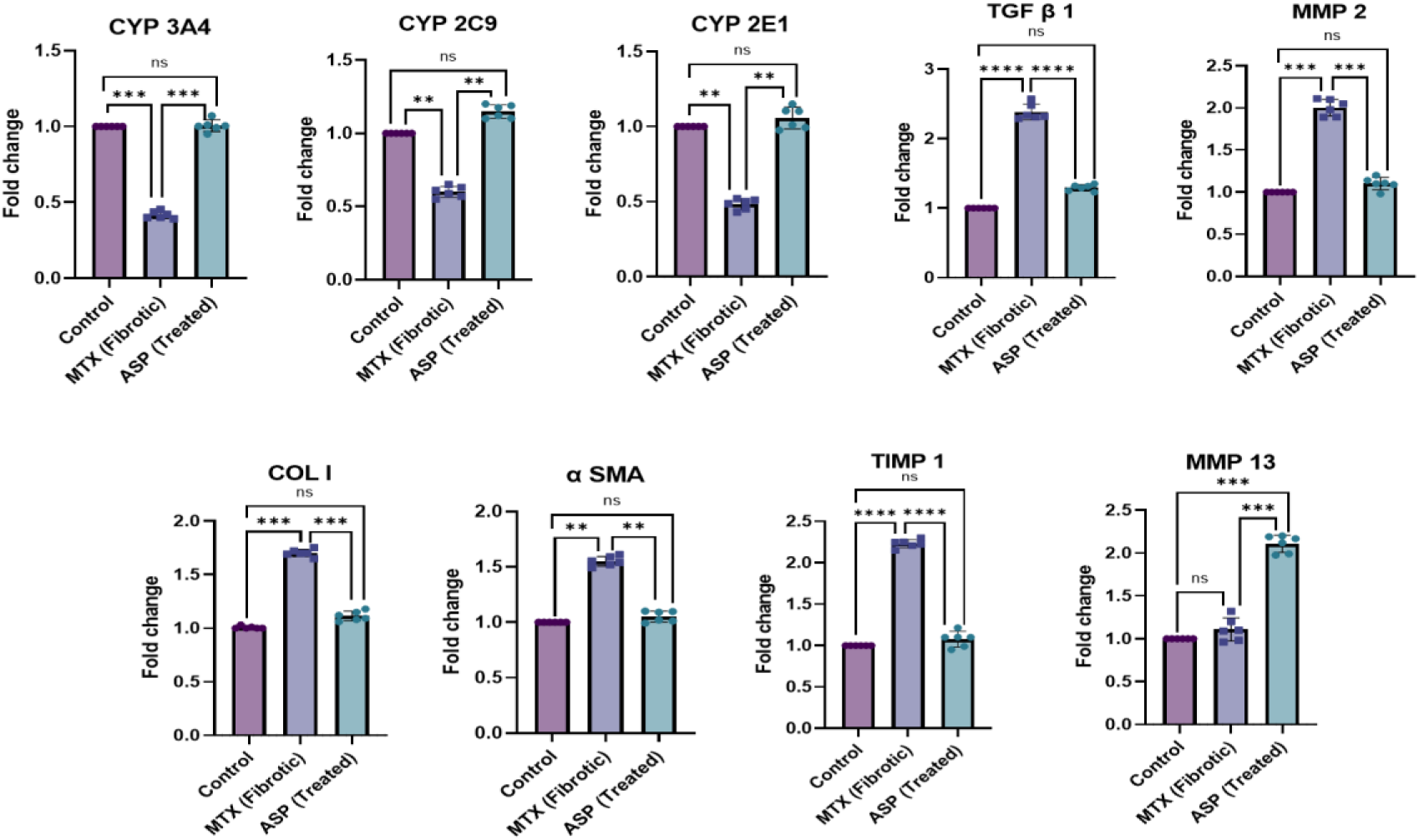
Gene expressions analysis performed on the 3D scaffolds; statistically significant ** : p<0.01, *** : p<0.05, **** : p< 0.0001 and ns: non-significant

Fibrosis-associated matrix. In contrast, collagen I (COL1A1) was markedly upregulated in MTX-fibrotic constructs, reflecting enhanced ECM deposition. ASP treatment reduced COL1A1 expression compared with the MTX group, trending back toward control levels.

Across targets, the directional changes lead loss of hepatocyte identity with MTX and its recovery with ASP, and ECM gene induction with MTX and its attenuation with ASP have mirror the histology (Section 3.3.3) and biochemical readouts (Section 3.3.4). Together, these transcript-level data substantiate that ASP reverses fibrotic programming while preserving hepatic function within the 3D GelMA–dECM model.

## 4.0 CONCLUSION

We developed a physiomimetic 3D GelMA–rat liver dECM bioprinted platform that recapitulates key structural and functional attributes of the human hepatic microenvironment and supports robust, controllable induction of fibrosis. MTX exposure produced hallmarks of fibrogenesis with elevated Ishak score, spindle-shaped/myofibroblast-like morphology, increased injury markers (LDH/ALT/ALP), suppression of hepatocyte function (albumin/urea secretion), and upregulation of COL1A1 with downregulation of ALB. Therapeutic challenge with aspirin (2 mM) reversed these phenotypes without compromising viability: histology regressed toward typical hepatic architecture (Ishak 4 → 1), albumin and urea rebounded while injury enzymes declined to control-comparable levels, and gene expression shifted toward a non-fibrotic program (↓COL1A1, ↑ALB). Collectively, these convergent readouts demonstrate that our 3D construct is a quantitative, human-relevant platform for evaluating anti-fibrotic candidates and mechanistic hypotheses, narrowing the translational gap between 2D screening and animal model. By coupling ECM fidelity with bioprinting precision, this platform enables ethically aligned, scalable R&D 3.0 workflows for phenotypic drug discovery, toxicity assessment, and mechanistic interrogation in liver fibrosis, with clear potential to accelerate preclinical down-selection of anti-fibrotic therapeutics.

## 5. ACKNOWLEDGEMENTS

The authors wish to acknowledge and thank the Department of Biotherapeutics Research (DBR), Manipal Academy of Higher Education (MAHE), and Manipal for the provision of research instruments and infrastructure support.

## 7. AUTHOR’S CONTRIBUTION

Kirthanashri S. V.: Conceptualization, Investigation, Data Curation, Writing – original draft, editing, reviewing and validation; Mrunmayi Gadre: Investigation, Data Curation, Visualization, Writing – original draft

## 8. ETHICS APPROVAL AND CONSENT TO PARTICIPATE

Not applicable.

## 9. CONSENT FOR PUBLICATION

Not applicable.

## 10. AVAILABILITY OF DATA AND MATERIALS

The datasets generated and/or analysed during the current study are available online as the Indian patent, repository (Indian Patent No. 202341045531)

